## Supplementary Material for "Evaluation of mitochondrial DNA copy number estimation techniques"

**Supplemental Table 1.** Picard Sequencing Summary Metrics Definitions

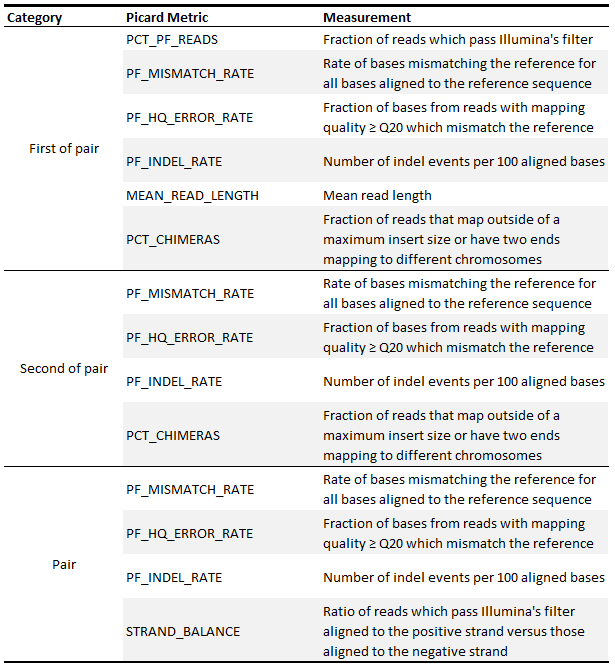

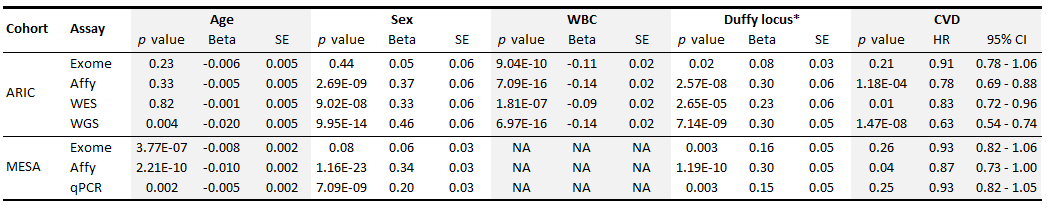

**Supplemental Table 2.** Associations of Known Correlates with mtDNA-CN Estimation Platforms

*Duffy locus associations were performed in blacks only

*Duffy locus associations were performed in blacks only

**Supplemental Table 3.** Relative performance of methods as rated by standardized –log *p* values

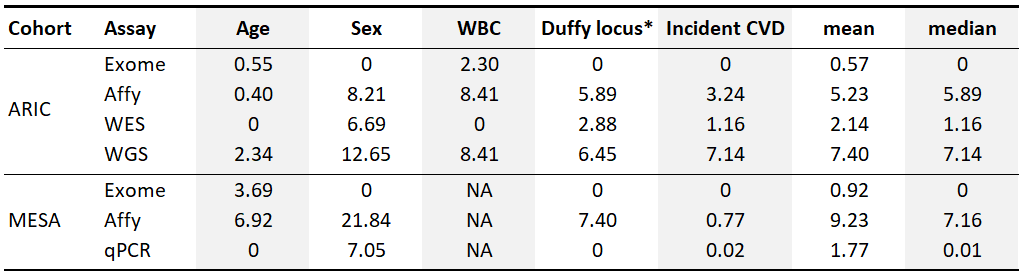

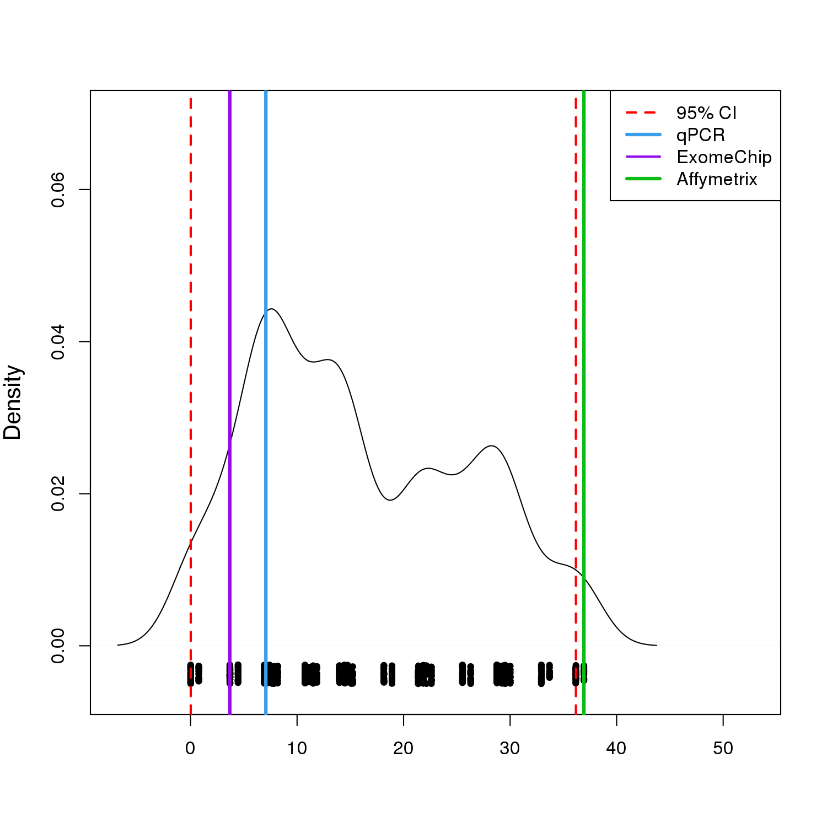

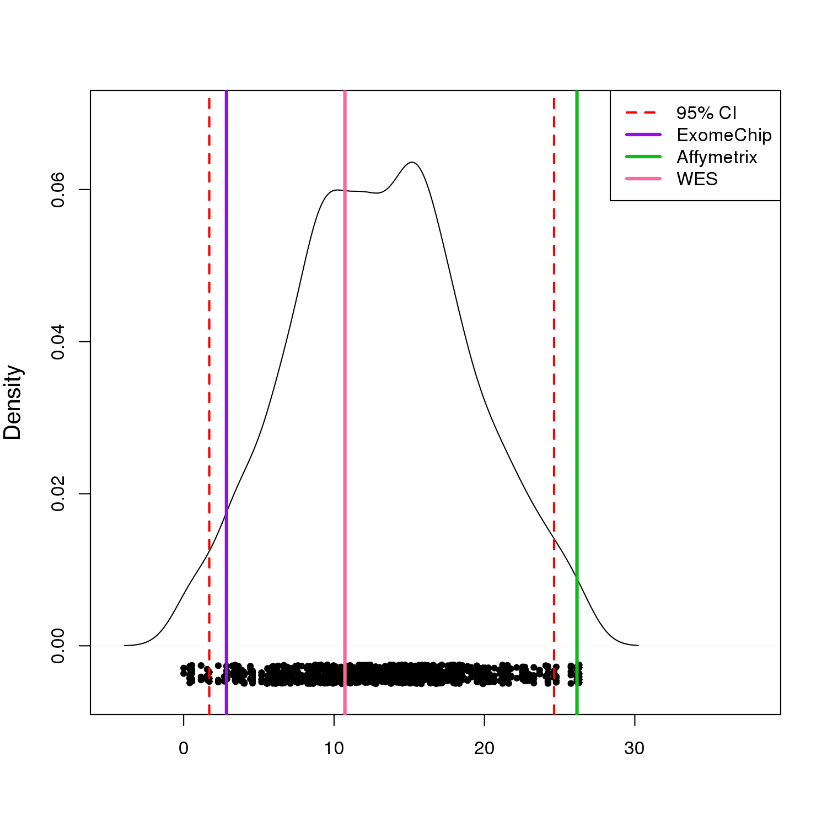

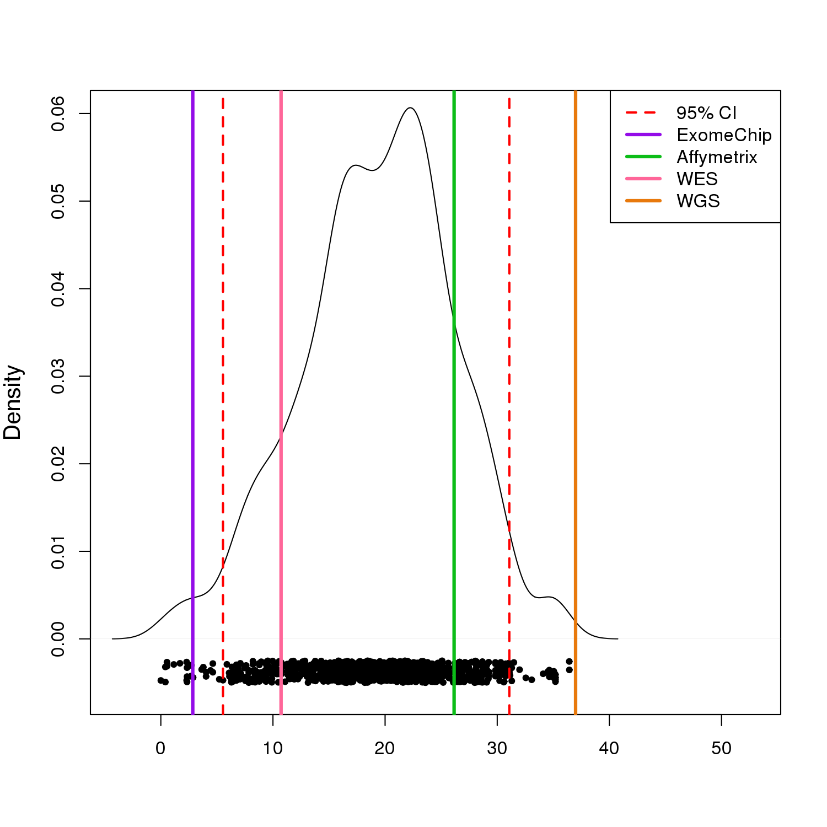

**Supplemental Figure 1. Permutation test for mtDNA-CN estimation method performance**

Performance scored as sum of negative log-transformed *p* value for each method across all correlates normalized to the least significant method of each correlate. Compared to 1,000 permutations of the sums of randomly selected normalized and transformed *p* values. ARIC (A), MESA (B), ARIC without WGS (C).

**A**

**B**

**C**

**Supplemental Table 4.** Relative performance of WGS and Affymetrix as rated by standardized –log *p* values

*Duffy locus associations were performed in blacks only

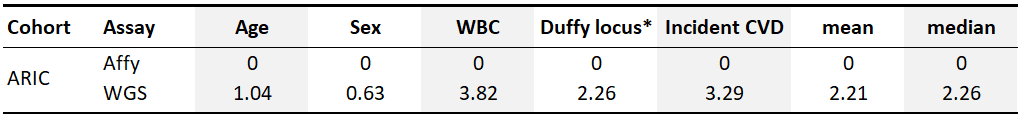

**A**

**B**

**Supplemental Figure 2**. **Phenotype Correlation plots**

ARIC (A) and MESA (B)

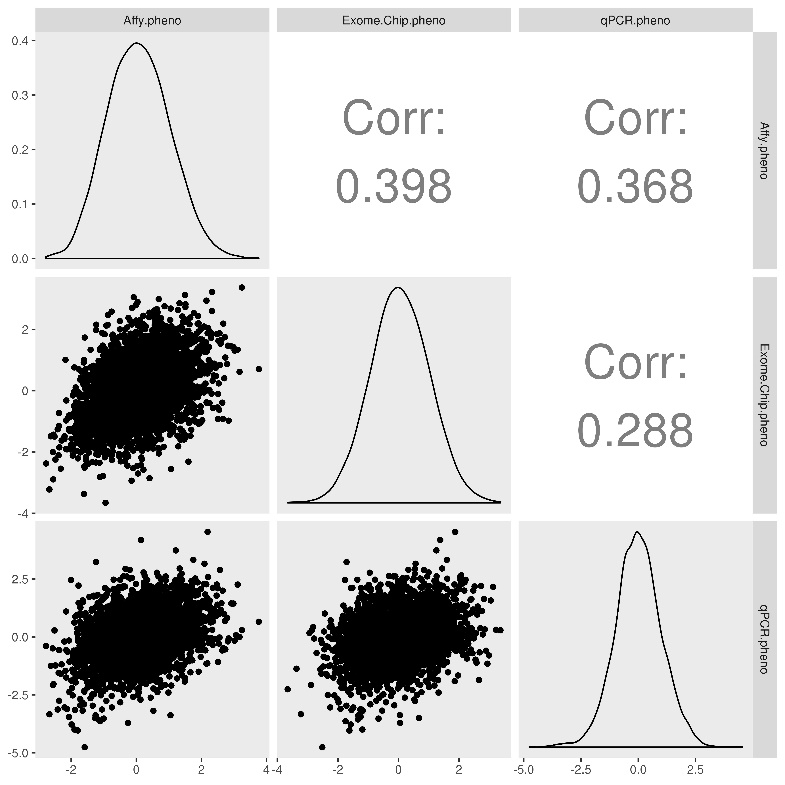

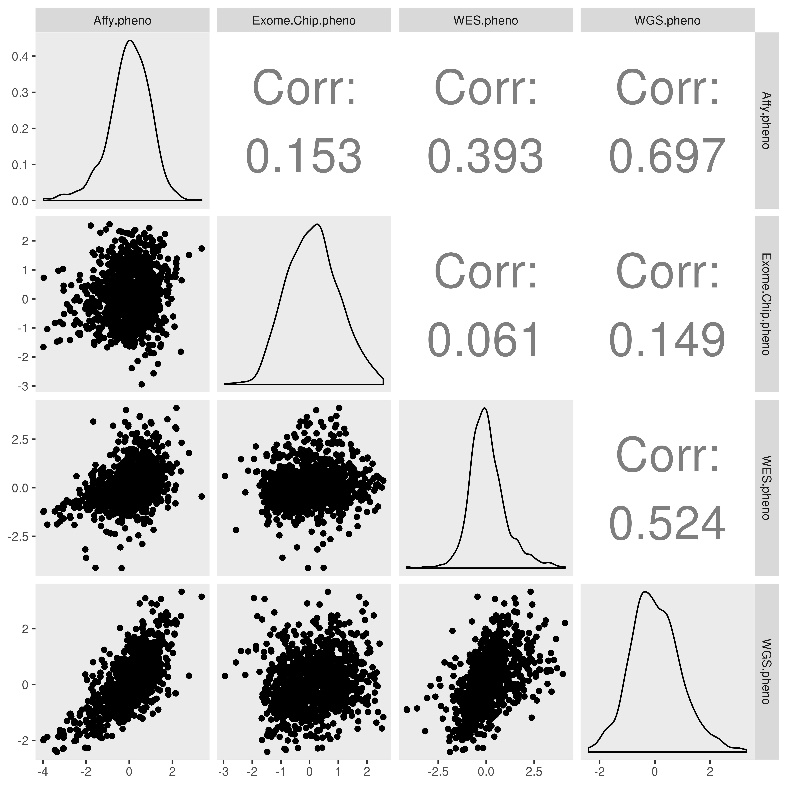

**Supplemental Table 5.** Participant Characteristics for dPCR Subset

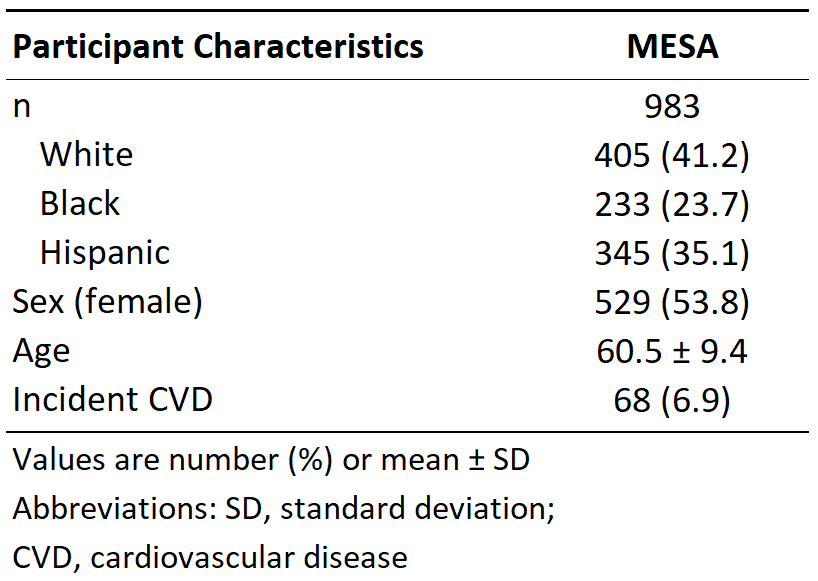

*Duffy locus associations were performed in blacks only

**Supplemental Table 6.** Associations of Known Correlates with mtDNA-CN Estimation Platforms for dPCR subset

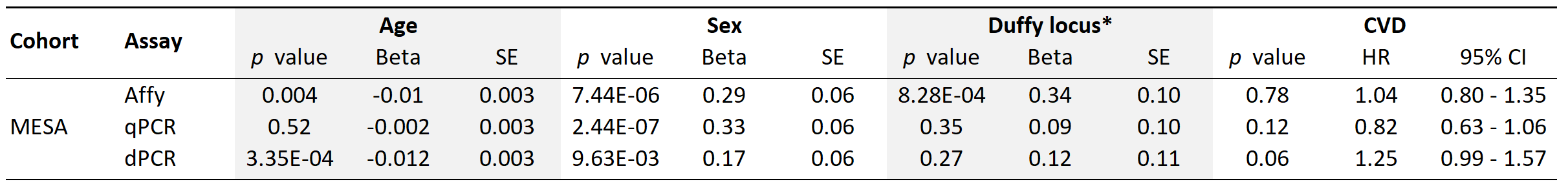

**Supplemental Table 7.** Relative performance of methods as rated by standardized –log *p* values for dPCR subset

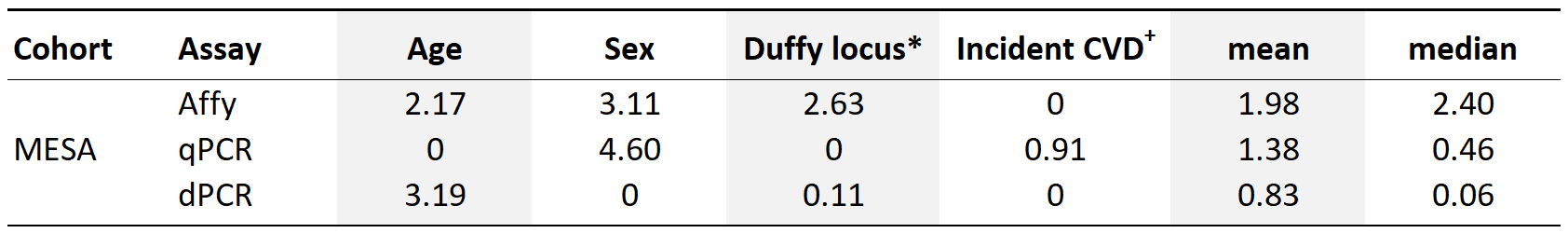

*Duffy locus associations were performed in blacks only

^+^Affymetrix and dPCR effect size estimates were in opposite direction as known effects and thus the -log *p* value of qPCR was standardized to a *p* value of 1 for Affymetrix and dPCR.
